# Restoring wild type-like network dynamics and behaviour during adulthood in a mouse model of schizophrenia

**DOI:** 10.1101/151795

**Authors:** Thomas Marissal, Rodrigo F. Salazar, Cristina Bertollini, Sophie Mutel, Mathias De Roo, Ivan Rodriguez, Dominique Müller, Alan Carleton

**Author notes:** These authors contributed equally to the work (Thomas Marissal, Rodrigo F. Salazar, Cristina Bertollini). Deceased on 29 April 2015 (Dominique Müller). Correspondence should be addressed to A.C. or T.M.

## Abstract

Schizophrenia is a severely debilitating neurodevelopmental disorder. Establishing a causal link between circuit dysfunction and particular behavioural traits relevant to schizophrenia is crucial to shed new light on the mechanisms underlying the pathology. Here we studied an animal model of the 22q11 deletion syndrome, which is the highest genetic risk to develop the pathology. We report a desynchronization of hippocampal neuronal assemblies that resulted from parvalbumin interneuron hypoexcitability. Rescuing parvalbumin interneuron excitability with pharmacological or chemogenetic approaches is sufficient to restore wild type-like network dynamics and behaviour during adulthood. In conclusion, our data provide mechanistic insights underlying network dysfunction relevant to schizophrenia and demonstrate the potential of reverse engineering in fostering new therapeutic strategies to alleviate the burden of neurodevelopmental disorders.

## Main text

Alterations of network dynamics have been proposed to be instrumental in schizophrenia^1^^-^^3^. A specific population of inhibitory neurons, the parvalbumin interneurons (PVIs), plays a key role in regulating network dynamics^4^^-^^7^ and may be involved in the pathology^8-11^. Although specific manipulations of PVI can reproduce behavioural phenotypes relevant to schizophrenia in rodents^12,13^, it remains unclear whether PVI dysfunction is causally linked to network dysfunction and pathological behaviour associated with schizophrenia. More importantly, it is not known whether manipulating PVI could restore altered physiology.

Among various genetic alterations, the specific deletion of ~30 genes on chromosome 22 that leads to the 22q11 deletion syndrome (22q11DS), is the highest identified genetic risk to develop schizophrenia^14,15^. We used a genetically engineered mouse bearing a hemizygous deletion on chromosome 16, termed Lgdel/+, which replicates the chromosomal alteration of the human 22q11DS^16^. In the CA1 area of the hippocampus, mouse models of 22q11DS differ from wild-type (WT) animals regarding their structural^17^^-^^19^ and electrophysiological properties^20^, and their functional connectivity with distant brain areas^3^. We first tested whether those differences were accompanied by intrinsic differences in network dynamics. Neural activity was monitored in hippocampal slices using the genetically encoded calcium indicator GCaMP6s expressed by CA1 neurons following adeno-associated viral (AAV) vector transfection (Fig. 1a,b). Network dynamics were induced by bath application of carbachol (50 μM), which triggered spontaneous calcium activity in individual neurons of wild type (WT) mice^21,22^ (Fig. 1c,d). Likewise, individual CA1 neurons of Lgdel/+ mice exhibited spontaneous calcium activity during the duration of the recording. Neither the fraction of active neurons (Fig. 1e), nor the mean frequency (Fig. 1f), the mean amplitude (Fig. 1g) and the mean duration (Fig. 1h) of calcium transients were significantly different between genotypes. In contrast, CA1 ensemble dynamics was strongly altered in Lgdel/+ mice. Indeed, neurons were largely desynchronized in respect to each other and the oscillations observed at a population level were largely reduced in Lgdel/+ mice (Fig. 1c,d). We thus quantified population activity and neuronal correlations and compared these quantities to data randomly shuffled in time. This enabled to control for the influence of neuronal co-activation occurring by chance (Supplementary Figs. 2 and 3). Lgdel/+ mice displayed less co-active cells (Fig. 1i), fewer occurrences of these co-activations (Fig. 1j) and less correlated neurons (Fig. 1k,l) in respect to WT animals. In summary, brain slices from Lgdel/+ mice exhibited a strong desynchronization of the CA1 hippocampal network.

**Figure 1.**
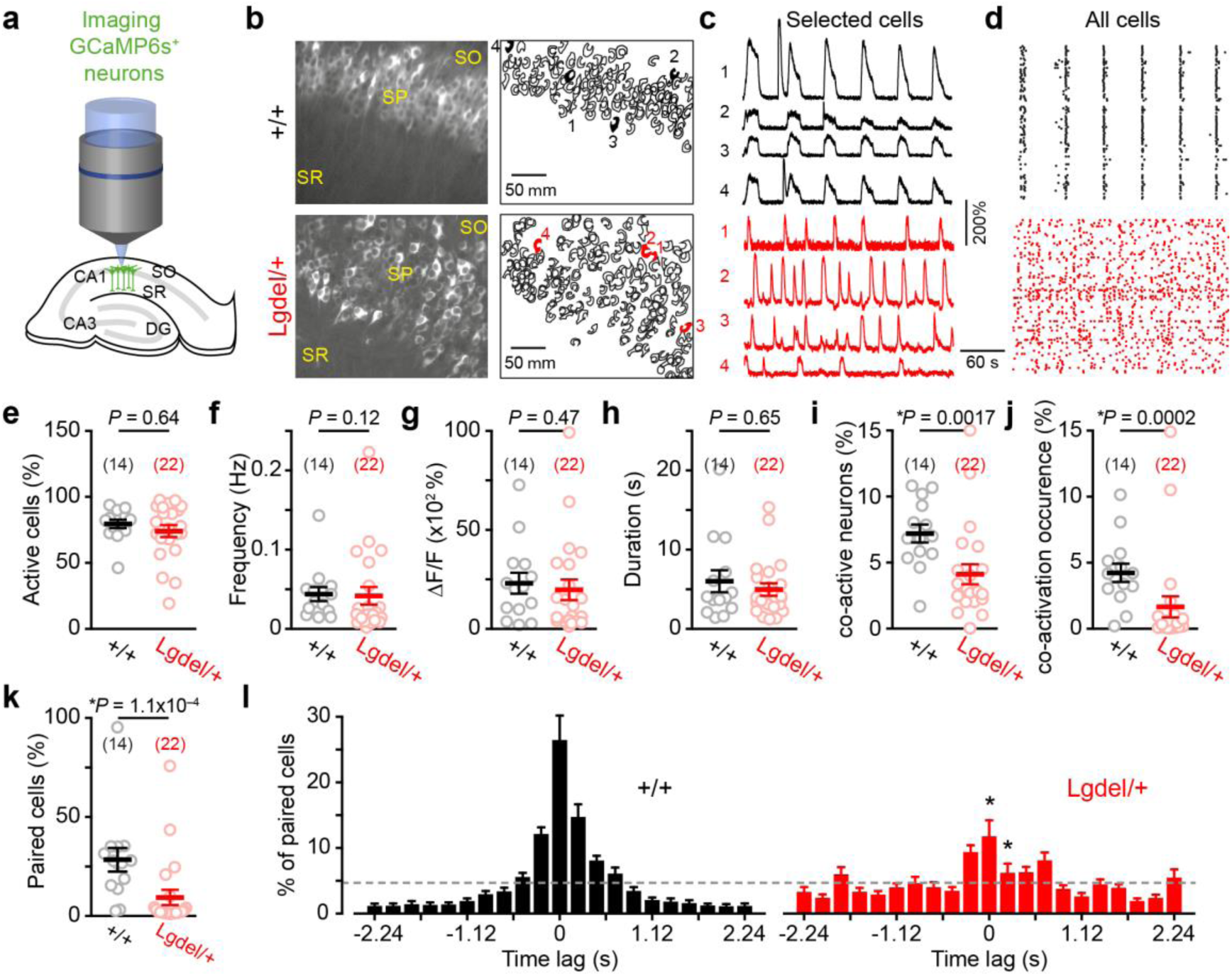
Desynchronization of CA1 neural assemblies in Lgdel/+ mice. (**a**) Schema of the experimental procedure. CA: Cornu Ammonis, DG: dentate gyrus, SO: stratum oriens, SP: stratum pyramidale, SR: stratum radiatum. (**b**) Photographs (*left*) and cell contours (*right*) of GCaMP6s-expressing cells in the CA1 region of WT and Lgdel/+ slices. (**c**) Examples of Ca^2+^ sweeps recorded from selected neurons identified as 1 to 4 in **b**. (**d**) Raster plots of Ca^2+^ transient onsets extracted from all neurons shown in **b**. (**e**) Proportion of neurons displaying Ca^2+^ transients in the CA1 region (each circle represents the mean for a given slice, number of slices indicated in parentheses). (**f-h**) Frequency (**f**), amplitude (**g**) and duration (**h**) of Ca^2+^ transients recorded in neurons of the CA1 area. The amplitude did not differ significantly despite a trend for a more prominent subpopulation of neurons with low firing rate in Lgdel/+ mice compared to WT mice (Supplementary Fig. 1a,b). (**i-j**) Percentage of co-active neurons above random co-activation (**i**) and occurrence of these co-activations (**j**). (**k**) Percentage of correlated pairs of neurons above chance. (**l**) Distributions of significant time lags in correlated pairs. The grey dash line represents the equidistribution. Data are presented as mean ± SEM. Statistical comparisons were done either with a Mann-Whitney test (**e-k**) or a two-way repeated measures ANOVA with Bonferroni post-hoc test (l): \**P* < 0.05.

To gain further insights into the mechanism underlying hippocampal desynchronization, we performed patch-clamp recordings of various neuronal populations in the CA1 region. First, we recorded spontaneous excitatory and inhibitory post-synaptic currents (sEPSCs and sIPSCs, respectively) in pyramidal cells (Fig. 2a). Consistent with a previous report^20^, CA1 pyramids recorded from Lgdel/+ mice had a comparable number of sEPSCs but less sIPSCs per second compared to CA1 pyramids in WT mice (Fig. 2a-d). In addition, no difference in sPSC amplitude (Supplementary Fig. 4a-c), in mEPSC frequency (Supplementary Fig. 4d) and in mIPSC frequency (Fig. 2e) was observed between the two genotypes. These results provide evidence that hippocampal pyramids in Lgdel/+ mice were characterized by hypoinhibition, likely to originate from the firing activity of GABAergic neurons.

**Figure 2.**
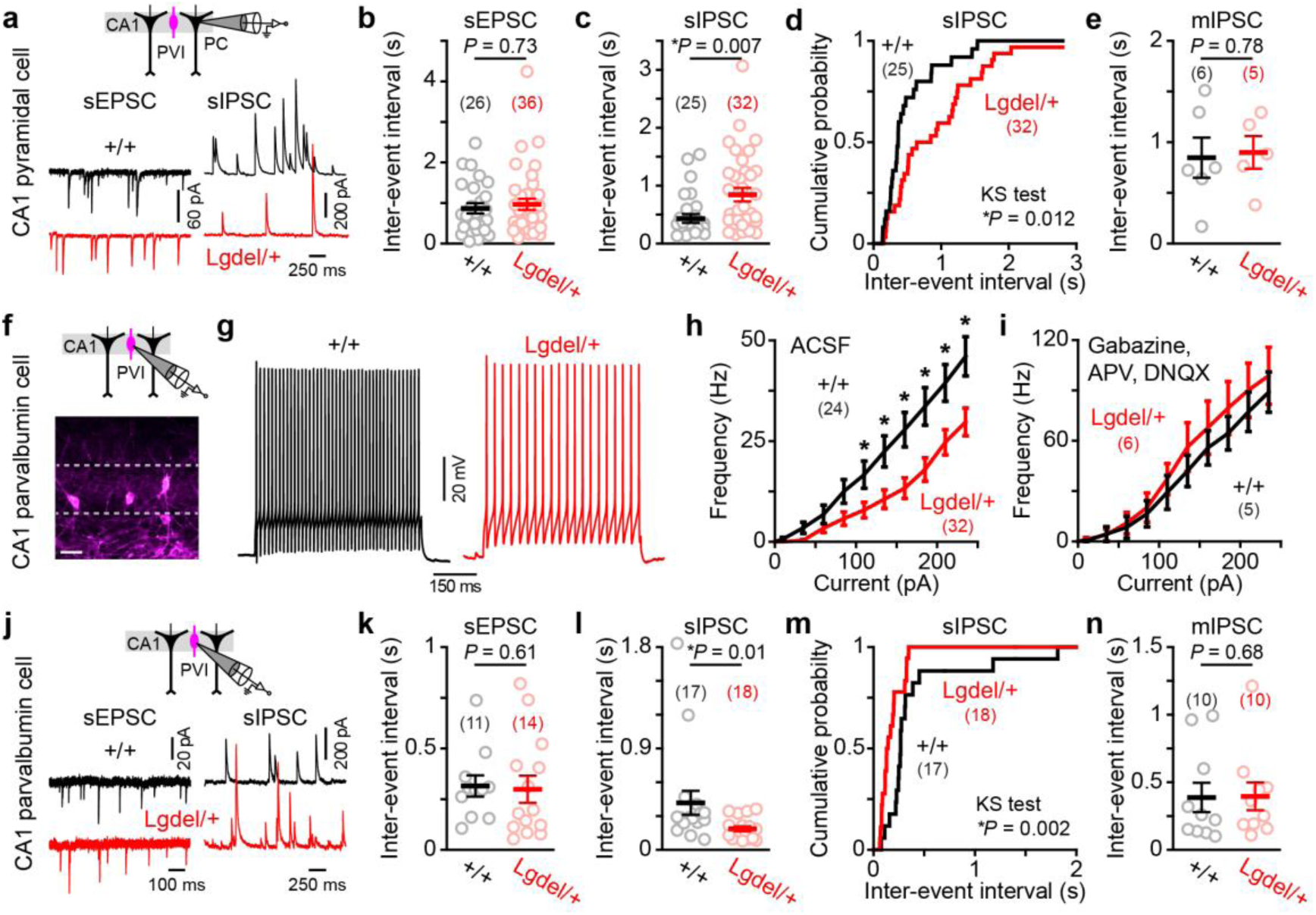
Reduced inhibitory inputs to CA1 pyramidal cells and hypoexcitability of CA1 parvalbumin interneurons in Lgdel/+ mice. (**a-e**) Patch-clamp recordings of CA1 pyramidal cells (PC) from WT and Lgdel/+ mice. (**a**) Examples of voltage-clamp recordings showing either spontaneous excitatory (sEPSC) or inhibitory (sIPSC) post-synaptic currents recorded from PC held at either −60 mV or 0 mV, respectively. (**b**) Summary graph of the mean sEPSC inter-event interval (IEI). (**c-d**) Summary graph (**c**) and cumulative distribution (**d**) of the mean sIPSC IEI (KS: Kolmogorov-Smirnov). (**e**) Summary graph of the mean miniature IPSC IEI. (**f-n**) Patch-clamp recordings from CA1 parvalbumin interneurons (PVIs) from *Pvalb*^cre/+^;+/+ (*black*) and *Pvalb*^cre/+^;Lgdel/+ (*red*) mice injected with AAV conditionally expressing the red fluorophore tdTomato. (**f**) Photograph of CA1 PVIs expressing tdTomato (scale bar: 50 μm). (**g**) Examples of spiking activity evoked by the same depolarizing step of current in CA1 PVIs recorded from WT and Lgdel/+ mice. (**h-i**) Input/output functions for CA1 PVIs recorded from WT and Lgdel/+ mice in either artificial cerebral spinal fluid (ACSF, i; difference between genotypes: two-way repeated measures ANOVA *F_1,9_* = 9.7 *P* = 0.003; *: post-hoc Fisher test at least *P* < 0.05) or in blockers of the fast glutamatergic and GABAergic synaptic transmission (**h**; two-way repeated measures ANOVA *F_1,9_* = 0.42 *P* = 0.54). (**j**) Examples of voltage-clamp recordings showing either sEPSC or sIPSC recorded from PVIs held at either −60 mV or 0 mV, respectively. (**k**) Summary graph of the mean sEPSC IEI. (**l-m**) Summary graph (**l**) and cumulative distribution (**m**) of the mean sIPSC IEI. (**n**) Summary graph of the mean mIPSC IEI. Data are presented as mean ± SEM (each circle represents a cell, number of cells indicated in parenthesis). Statistical comparisons were done using a Mann-Whitney test unless otherwise stated.

We further tested the latter hypothesis by recording genetically labelled PVIs (Fig. 2f), which exert an inhibitory control over pyramidal cells^23,24^ in the hippocampal region. An analysis of input/output function in current-clamp experiments revealed that CA1 PVIs were less excitable in Lgdel/+ than in WT animals when recorded in artificial cerebrospinal fluid (Fig. 2g,h). Interestingly, such firing difference disappeared when GABAA, AMPA and NMDA receptors were blocked pharmacologically (Fig. 2i). Thus, the PVI hypoexcitability may rather reflect differences in PVI synaptic inputs than changes of their intrinsic properties. Supporting this notion, CA1 PVIs recorded from Lgdel/+ mice received similar number of sEPSCs but more sIPSCs per second than CA1 PVI neurons recorded in WT animals (Fig. 2j,k). No differences in sPSC amplitude (Supplementary Fig. 4G-I), mEPSC frequency (Supplementary Fig. 4j) and mIPSC frequency (Fig. 2n) were observed. Taken together, our data show that CA1 pyramidal cells of Lgdel/+ mice are hypoinhibited and that PVI are hypoexcitable.

Could network desynchronization be due to PVI hypoexcitability? We used a cell-autonomous chemogenetic approach by specifically infecting CA1 PVIs from *Pvalb*^cre/+^;+/+ mice with AAV enabling the conditional expression of the inhibitory designer receptor exclusively activated by designer drug (DREADD) hM4Gi, which is selectively activated by the inert drug clozapine-N-oxide (CNO)^25^. Effectively, reducing PVI excitability desynchronized CA1 assemblies in WT mice (Supplementary Fig. 5). Thus, could hippocampal network dynamics be rescued in Lgdel/+ mice by increasing PVI excitability? First, we tested a pharmacological approach by using a Neuregulin 1 peptide (NRG1), which increases PVI excitability in WT mice^26,27^. Second, we specifically expressed the excitatory DREADD hM3Gq in CA1 PVIs from *Pvalb*^cre/+^;Lgdel/+ mice. During path-clamp recordings of hippocampal slices, both strategies (Fig. 3a-f) raised PVI excitability and restored input/output functions comparable to the ones recorded from WT mice (difference between WT and treatments in Lgdel/+: two-way repeated measures ANOVA *F_1,29_* = 0.44, *P* = 0.51 and *F_1,28_* = 0.12, *P* = 0.73 for NRG1 and CNO treatment, respectively). We then tested the effect of the two treatments on CA1 network dynamics by pre-incubating hippocampal slices with either NRG1 or CNO prior to calcium imaging. Strikingly, both strategies increased neuronal correlations (Fig. 3g-j) and co-activations (Supplementary Fig. 6) in Lgdel/+ mice to a level comparable to WT littermates. Thus, counterbalancing the PVI hypoexcitability of Lgdel/+ mice was sufficient to extinguish the network desynchronization observed in brain slices.

**Figure 3.**
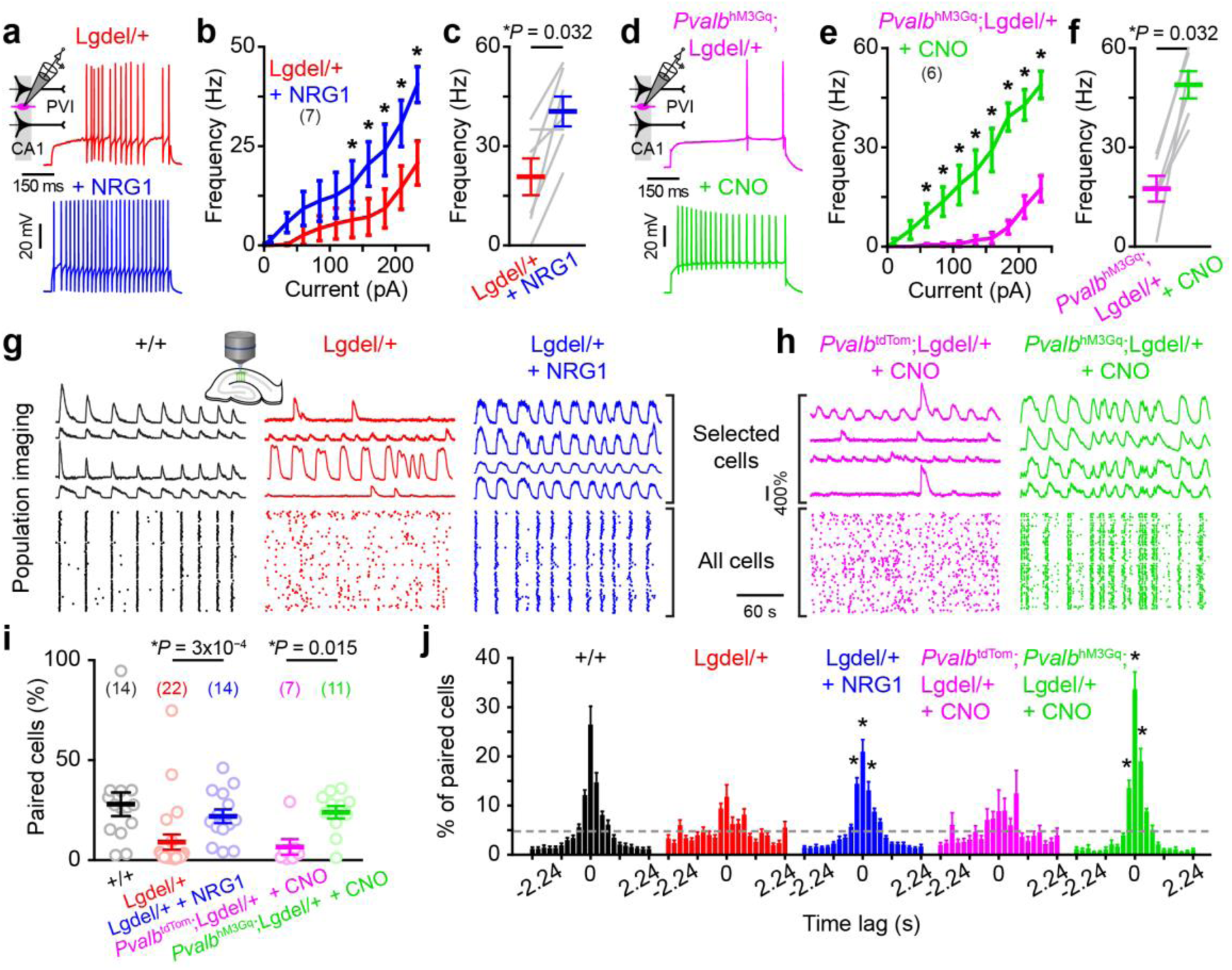
Rescue of PVI excitability restores synchronization of CA1 assemblies. (**a**) Examples of spiking activity evoked by the same depolarizing step of current in CA1 PVI recorded from *Pvalb*^cre/+^;Lgdel/+ mice before and after neuregulin 1 (NRG1, 10 nM) bath application. (**b**) Input/output functions for CA1 PVI recorded before and after NRG1 application (drug effect: two-way mixed design repeated measures ANOVA *F_1,6_* = 11.4 *P* = 0.015; *: post-hoc Fisher test at least *P* < 0.05). (**c**) Summary graph of the firing frequency generated by PVIs in response to a 235 pA current step before and after NRG1 application (Wilcoxon paired test). (**d**) Examples of spiking activity evoked by the same depolarizing step of current in CA1 PVI recorded from *Pvalb*^cre/+^;Lgdel/+ expressing the excitatory DREADD hM3Gq (*Pvalb*^hM3Gq^;Lgdel/+) before and after clozapine-N-oxide (CNO, 1 μM) bath application. (**e**) Input/output functions for CA1 PVIs recorded before and after CNO application (Drug effect: two-way mixed design repeated measures ANOVA *F_1,5_* = 55.2 *P* = 7x10^−***4***^; *: post-hoc Fisher test at least *P* < 0.05). (**f**) Summary graph of the firing frequency generated by PVI in response to a 235 pA current step before and after CNO application (Wilcoxon paired test). (**g-h**) Examples of Ca^2+^ sweeps recorded from selected neurons and of raster plots of Ca^2+^ transient onsets extracted from all imaged neurons in slices from various genotypes and following various drugs application. (**i**) Percentage of correlated pairs of neurons above chance (each circle represents a given slice, number of slices indicated in parentheses; Mann-Whitney test). (**j**) Distributions of the time lag computed for correlated pairs. The grey dash lines represent the equidistribution (two-way repeated measures ANOVA with Bonferroni post-hoc \**P* < 0.05). Data are presented as mean ± SEM.

Although 22q11DS is considered to be a neurodevelopmental disorder, we investigated whether the same pharmacological and chemogenetic treatments are efficient in adult animals. We first performed *in vivo* electrophysiological recordings in the dorsal CA1 area (dCA1) of awake mice (Fig. 4a and Supplementary Fig. 7a). A spectral analysis of the local field potential (LFP) revealed lower power in Lgdel/+ mice compared to WT animals in the theta frequency band (5-8Hz; Fig. 4a,b and Supplementary Fig. 7b, Kolmogorov-Smirnov test *P* = 8.1 x 10^−4^). Neuronal oscillations in the theta frequency band are crucial to hippocampal functions^28^ and their strength can be modulated by PVIs activations^5^. Interestingly, NRG1 injections increased the power of theta oscillations to levels close to WT mice (Fig. 4a-c, Kolmogorov-Smirnov test *P* = 0.1). To improve the specificity of our manipulation, we infected dCA1 PVI of *Pvalb*^cre/+^;Lgdel/+ mice with an AAV enabling the conditional expression of the hM3Gq DREADD (Supplementary Fig. 7c). CNO injections also increased the LFP power up to levels comparable to WT mice (Fig. 4a-c, Kolmogorov-Smirnov test *P* = 0.67). In summary, NRG1 injection and chemogenetic excitation of PVIs in adult animals were sufficient to establish WT-like network dynamics in awake Lgdel/+ mice.

**Figure 4.**
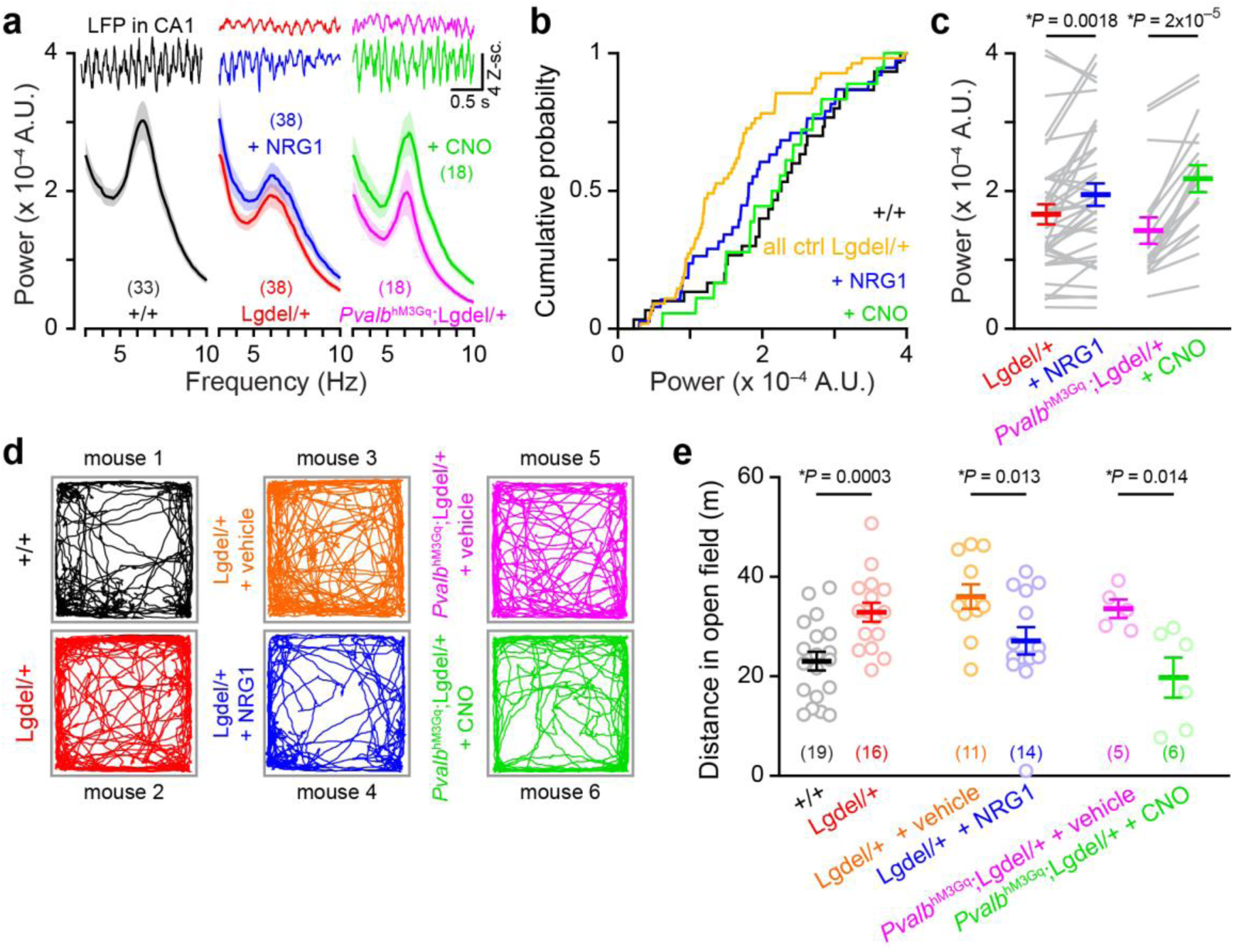
Increasing PVI excitability during adulthood restores hippocampal theta oscillations and behaviour. (**a**) Representative sweeps of local field potentials (LFPs) recorded in the CA1 region of awake mice. The distributions of LFP power spectra are presented (mean ± SEM; number in parentheses indicate the number of shank/mouse pairs) for WT, Lgdel/+ before and after NRG1 injection, and *Pvalb*^hM3Gq^;Lgdel/+ mice before and after CNO induced effect (see methods). (**b**) Cumulative probabilities for the different transgenic mice and treatments. As the baselines values of LFP power (averaged between 5 to 8 Hz for the theta frequency band) of Lgdel/+ mice and *Pvalb*^cre/+^;Lgdel/+ did not differ statistically (Kolmogorov-Smirnov KS test *P* = 0.2), they were grouped and referred to as Lgdel/+ control mice (‘all ctrl Lgdel/+‘). See main text for statistics. (**c**) Power in the theta frequency range in the control conditions or after treatments with NRG1 (*left*) or CNO (*right*), respectively (Wilcoxon paired test). (**d**) Locomotor tracks of different transgenic mice and treatments (mouse 1 to 6) during 10 min in an open field. (**e**) Summary graph of the distance travelled in the open field (each circle represents a mouse; number of mice analysed indicated in parentheses; Mann-Whitney test). Data are presented as mean ± SEM.

Could the same pharmacological and chemogenetic manipulations also affect hippocampal-dependent behaviours? As observed in various animal models of schizophrenia^29^ including the 22q11DS mouse model^18^, adult Lgdel/+ mice exhibit hyper-locomotor activity in comparison to their WT littermates in the open field test (Fig. 4d,e). Intraperitoneal injections of NRG1 in Lgdel/+ mice (30-60 min prior to open field testing) and selective excitation of dCA1 PVI expressing the hM3D DREADD led to a reduction in the distance travelled to levels similar to the ones reached by WT littermates (Fig. 4d,e; Mann-Whitney test *P* = 0.15 and *P* = 0.55, respectively). Therefore increasing PVI excitability in adult Lgdel/+ mice was sufficient to induce a behavioural pattern similar to WT littermates, though the converse is not true. Indeed, when we selectively inhibited dCA1 PVI expressing the hM4D DREADD, the locomotion of WT mice was not significantly affected^12^ (Supplementary Fig. 8a,b).

In conclusion, our findings provide a mechanistic explanation of the alterations observed in an animal model of 22q11DS at the cellular and network levels. Our work suggests that the hypo-excitability of inhibitory neurons such as PVI leads to an alteration of the neuronal synchronization in the hippocampus and probably in other brain areas^30^. Although the 22q11DS is a neurodevelopmental disorder, our data demonstrate that selective neuronal manipulations during adulthood are sufficient to restore functional brain dynamics and typical behavioural patterns. Furthermore, our results highlight PVIs as a valuable therapeutical target for 22q11DS and similar neuropsychiatric disorders.

## AUTHOR CONTRIBUTIONS

T.M., D.M. and A.C. carried out the study conceptualization. T.M., R.S., C.B., S.M., M.D.R., I.R., D.M. and A.C. contributed to the experimental design. TM performed the calcium imaging experiments and their analyses. C.B. carried out the *in vitro* electrophysiological experiments. R.S. performed the *in vivo* electrophysiological recordings and developed most of the MATLAB-based programs used for the analysis of the calcium imaging and the electrophysiological recordings. M.D.R. developed some MATLAB-based scripts used to analyze calcium imaging data. T.M. performed behavioral experiments with the help of S.M. A.C., T.M., R.S. and I.R. wrote and edited the manuscript with comments from all other authors.

## ACKNOWLEDGEMENTS

This paper is dedicated to the memory of Dominique Müller, whose ideas inspired this study and will continue to inspire all of us. We thank Raju Kucherlapati for generously providing the Lgdel/+ mice. We thank Lorena Jourdain and MariePriscille Hervé, for their technical support. We thank Yann Bernardelli, Pablo Mendez Garcia, Thomas Stefanelli, Stéphane Eliez, Pico Caroni and other members of the NCCR SYNAPSY for helpful discussions and/or comments on the manuscript. This research was supported by the University of Geneva, the Swiss National Science Foundation (grant numbers: 31003A_172878 to A.C., 310030B_144080 to D.M. and 310030E_135910 to I.R.), the National Center of Competence in Research (NCCR) “SYNAPSY - The Synaptic Bases of Mental Diseases” financed by the Swiss National Science Foundation (grant 51NF40-158776, D.M. and A.C.) and the Lejeune Foundation (T.M.).

## SUPPLEMENTARY MATERIALS

### MATERIALS AND METHODS

#### Animals

Mice were maintained on a normal 12h light/dark cycle at 24°C with *ad libitum* access to food and water. We crossed Lgdel/+ (Del(16Dgcr2-Hira)1Rak ^16^, generously provided by Prof. Raju Kucherlapati, Harvard university), deleted from the 22q11DS-related genes, with C57BL/6J wild-type mice (Janvier, France) to generate Lgdel/+ mice and wild-type (WT) littermates. Homozygous *Pvalb*^cre/cre^ mice (Pvalb tm1(cre)Arbr/J ^31^, Jackson laboratory, #017320; C57BL/6J background) were crossed with Lgdel/+ mice to generate *Pvalb*^cre/+^;+/+ and *Pvalb*^cre/+^;Lgdel/+ littermates.

All animal protocols are in accordance with the Swiss Federal Act on Animal Protection, with the Swiss Animal Protection Ordinance and were approved by the University of Geneva and Geneva state ethics committees (authorization numbers: 1007/3129/0 and GE/156/14).

#### Adeno-associated viruses

Adeno-associated viruses (AAVs) were produced by the University of North Carolina and the University of Pennsylvania vector cores. Calcium signals were detected using a recombinant AAV1.Syn.GCaMP6s.WPRE.SV40, which enables the expression of the calcium-sensor GCaMP6s in neurons. The recombinant AAV5-hSyn-DIO-hM4D(Gi)-mCherry and AAV5-hSyn-DIO-hM3D(Gq)-mCherry construct were used for chemogenetic inhibition and excitation of Cre-expressing PVIs, respectively ^32^. AAV9.CAG.Flex.tdTomato.WPRE.bGH was used in order to specifically label Cre-expressing PVIs with a red fluorophore.

#### Slice preparation

Five to six day-old pups were decapitated and their hippocampi were carefully removed and unfolded. Transverse hippocampal slices (400 μm thickness) were made using a choper (McIlwain). Slices from dorsal and ventral parts of the hippocampus were used undistinguishable throughout the study. Slices were then maintained in an incubator at 33°C in a culture medium as previously described ^33^. Viral infection of the slices was performed four days after the preparation of the culture (day in vitro 4, DIV4), by putting a hydrophilic membrane (FHLC white membrane, Millipore) on the slice surface. A drop of AAV-containing solution (approximately 0.1 mL) was placed on the surface of the targeted CA1 region of the hippocampus using a pressurized glass pipette (Harvard Apparatus and Toohey Spritzer, Toohey Company). The membrane was left on the slice 1h in an incubator at 33°C before being removed. Calcium imaging and patch-clamp experiments were conducted around one week after the viral infection that is from DIV11 onwards.

#### Calcium imaging and analyses

Slices previously infected at DIV4 with an AAV virus leading to the expression of the genetically encoded calcium sensor GCaMP6s ^34^ under the control of the human synapsin promoter were used. In some experiments, AAV enabling the conditional expression of the excitatory DREADD hM3Gq or the red fluorophore tdTomato were also injected. Slices were immerged in an artificial cerebro-spinal fluid (ACSF) containing (in mM): NaCl (124), KCl (1.6), KH_2_PO_4_ (1.2), MgCl_2_ (1.3), CaCl_2_ (2) Glucose (10), and ascorbic acid (2.0), pH 7.4, continuously oxygenated and perfused using a peristaltic pump. Calcium transients were recorded using a Nipkow-type spinning disk confocal microscope (Olympus) coupled with single-photon lasers (excitation wavelength 488 and 565nm). Images were acquired through a CCD camera (Visitron Systems Evolve). Slices were imaged using a 10X 0.30NA objective (Olympus) at 8.9 Hz frame rate (i.e. 112 ms per frame). A few calcium-imaging experiments were performed (same frame rate as above) using a multibeam two-photon laser scanning system (Trimscope, LaVision Biotec, Germany) mounted on a BX51WI Olympus microscope coupled to Ultra II chameleon laser (Coherent). A typical imaging session covered a field of 420x420μm size containing ~250 individual neurons in the CA1 area. Slices were recorded for 6 minutes either before or five minutes after carbachol (CCh) application. For some experiments, slices were pre-incubated with NRG1 (10 nM) or CNO (1 μM) 15 minutes prior to CCh application of. All drugs remained present during the entire recordings.

Calcium data ^35^ analyses were performed using the Matlab-based ‘Caltracer3beta’ software (Columbia University), which enables a) the tracing of Regions Of Interests (ROIs) matching the identified GCaMP6S-expressing neuronal soma, b) the calculation of the averaged fluorescence signal from each cell-based ROIs as a function of time. In order to distinguish the actual neuronal calcium activity from background activity (i.e. fibers nearby cells), the fluorescence within a halo of the pixels surrounding the ROI was subtracted from the fluorescence signal recorded in the ROI. From the resulting signal, the onsets and offsets of the calcium events were identified. Onsets were automatically detected when fluorescence exceeded a threshold set for every slice (based on the background noise) during a minimum time of 1s (based on the kinetics of GCaMP6s), and offsets were defined as the events half-decay time. Onsets and offsets were manually corrected after automatic detection, and used to estimate 1) the proportion of active neurons and the frequency of calcium events, 2) the amplitude and the duration of calcium events, 3) the proportion of co-activated neurons and the duration of their co-activation and 4) the proportion of correlated pairs of neurons and their preferred lags. These analyses were performed using custom-made Matlab-based scripts.

Calcium event amplitude was calculated considering the cell fluorescence value at the onset of the event (F0) and the maximum value of fluorescence measured during the event (F), and was scored as (F-F0)/F0 in %. All signals below 5% were considered as noise. Calcium event duration was calculated by subtracting the offset to the onset time values. An average event amplitude and duration was calculated for each neuron in each experiment (point 2 above).

The co-activation of neurons (point 3 above) is the number of onsets over the whole population for each frame. This co-activation reached significance if its value was above the maximum number of onsets found in a surrogate population obtained by randomization. The randomization (10000 iterations) consisted in adding a random shift (between one and the total number of frames minus one) to the onsets of each neuron; each neuron had a different shift assigned. When the shifted onsets exceeded the last frame, the difference was referenced to the first frame. This procedure maintained the same activity pattern for each neuron but scrambled the relationship between neurons. This method provided an estimate of the fraction of co-active neurons that exceeded chance levels and the fraction of frames when it occurred.

The correlation between two neurons (point 4.) was quantified as the number of onset coincidences occurring between lags spanning -20 to +20 frames and binned into two frames. Significant correlation between two neurons occurred when the number of coincidences exceeded a chance threshold at any lag. The threshold was obtained as the maximum number of coincidences found in a surrogate population (2000 iterations) generated by a similar randomization procedure as described in the previous paragraph. The only difference was that the random shift occurred within the range of the lags used with the exclusion of zero-lag. This procedure highlights pairs of neurons that are co-active within 20 frames and that have preferred lags, usually only one, above chance.

#### Patch-clamp electrophysiology

Recordings were done from slices perfused with oxygenated ACSF (see composition above). CA1 pyramidal cells were visually identified using infrared video microscopy. PVIs were identified by fluorescence after the infection of *Pvalb*^cre/+^;+/+ and *Pvalb*^cre/+^;Lgdel/+ slices with AAVs (see above). For recordings in current clamp mode, cells were recorded using patch-clamp pipettes (tip resistance, 3–4 MΩ) filled with an intracellular solution containing (in mM) 70 K-gluconate, 70 KCl, 2 NaCl, 2 MgCl2, 10 HEPES, 1 EGTA, 2 MgATP, and 0.3 Na2GTP (pH 7.3) corrected with KOH (290 mOsm). For recordings performed in voltage-clamp mode, the intracellular solution contained the following (in mM): 125 CsMeSO3, 2 CsCl, 10 HEPES, 5 EGTA, 2 MgCl2, and 4 MgATP. Signals were amplified using a Multiclamp 200B patch-clamp amplifier (Molecular Devices), filtered and digitized at 4 kHz with an analog-to-digital converter (Digidata 1322A, Axon Instruments) and stored using pClamp 9 software (Axon Instruments). The analysis of electrophysiological recordings was performed using Clampfit 10.2 (Molecular Devices) and MiniAnalysis software (Synaptsoft).

Passive membrane properties were obtained immediately after membrane rupture. Series and membrane resistances were monitored continuously during voltage-clamp experiments by applying a -5 mV step at the beginning of every sweeps and recordings were discarded when these parameters changed by >20%. To record spontaneous and miniature inhibitory and excitatory post-synaptic currents (sIPSCs, mIPSCs, sEPSCs and mEPSCs, respectively), cells were held at 0mV and -60 mV, respectively. These values were chosen accordingly to the measured reversion potentials of glutamatergic and GABAergic currents. Miniature currents were recorded in the presence of TTX (1μM). Firing pattern were assessed in current clamp mode by injecting current in 25 pA steps of 600 ms of duration starting from -50 pA to 225 pA. The firing frequency was determined at each step. For some experiments, PVIs were recorded before and 15min after treatment with NRG1 (10 nM) or CNO (1 μM), or a mix of APV (50 μM), DNQX (10 μM) and Gabazine (10 μM).

#### Surgery

Analgesic treatment (paracetamol 0.2 g/kg) was administered for two days prior to the surgery. Anaesthesia was induced at 5.0% and maintained at 1.5%– 2.0% isoflurane (w/v) (Baxter AG). Mice were placed in a stereotaxic apparatus (Angle One) above a heating pad. Eyes were covered with ointment (Lacyvisc, Alcon) and Lidocaïne, a local analgesic, was injected below the skin overlaying the skull (s.c.). After exposing the skull, a craniotomy (3mm in diameter using a biopsy punch from Miltex) was made above the hippocampus. In *Pvalb*^cre/+^;Lgdel/+ mice, AAV infections (0.1 μl at 0.05 ml/min) were performed using glass micropipettes (10–15 μm tip diameter; Drummond Scientific Company) at AP: –1.7 mm, ML: 1.0 mm bilaterally and DV: 1.5 mm relative to Bregma and the skull surface in order to target the dorsal CA1 area of the hippocampus. A few infections (0.5 μl) were carried out using the coordinates AP: –1.5 mm, ML: 1.5 mm bilaterally and DV: 1.5 mm relative to Bregma and the skull surface. The micropipettes were left in place for 5 min after microinjection and slowly retracted (0.4 mm/min) to avoid reflux of the viral solution. In all mice used for in-vivo electrophysiological recordings, the craniotomy was covered by a glass cover slip (Ø 3mm, Warner), which was then glued (Cyanocrylate, Cyberbond) to the skull. A ground wire was positioned a couple of millimetres posterior to the recording craniotomy at the surface of the cortex. A head-post, enabling head restraining during electrophysiological recordings ^36,37^, was cemented to the skull using a mixture of dental cement (Jet Repair, Lang) and glue (Cyanocrylate, Cyberbond). AAV-Injected mice were used either for behavioural or electrophysiological measurements approximately three weeks after infection. All mice used for electrophysiological recordings had at least three days for recovery and three days of head restraining habituation. After the completion of each *in vivo* experiment, mice were injected i.p. with a lethal dose of pentobarbital (150 mg/kg) followed by intracardiac perfusion with 4% paraformaldehyde (PFA). Brains were maintained in 4% PFA for one day and stored in PBS for histological verification of the polytrode position and/or infection accuracy (Supplementary Fig. 7).

#### In vivo electrophysiological recordings and analyses

Animals were prepared with a subcutaneous catheter (if planned for s.c. injections) and then fixed to a head-restraining device as previously described ^38,39^. The cover slip protecting the brain was then taken out and the dura-matter removed under 1.5% isoflurane anaesthesia. After stopping the anaesthesia flux, a DiI (Thermo Fisher Scientific) covered polytrode (Buszaki 64L; Neuronexustech, USA) was slowly lowered (~15 min) between 1.2 and 1.7 mm DV from the surface of the cortex. Then, a period dedicated to the stabilization of the signal lasted between 10 to 20 min. Broadband signals (0.1-9000 Hz) were acquired at 32 KHz sampling rate (Neuralynx Co). The first experiment consisted in recording a baseline period of 30 min followed by a s.c. injection of Neuregulin (NRG1, 50 μg/kg) and a additional one hour period of drug induced neuronal activity. The second experiment consisted in injecting clozapine N-oxide (CNO, 2mg/kg) i.p. before setting up the animals in the head-restraining device. The periods considered as control and as CNO-mediated excitation effect (magenta and green conditions in Fig. 4a-c, respectively) consisted in 30-60 and 60-90 minutes post-injection, respectively. In both experiments, recordings were done for 1.5 hour. After completion of the experiments, the polytrode was slowly retracted and the animals were removed from the head-restraining holders.

Using the Matlab environment (MathWorks Co), local field potentials (LFP) were extracted by lowpass filtering (1-90 Hz) and downsampling (200Hz) broadband signals. After a linear detrend of the signal, the LFP for each channel was normalized using z-scores. The spectral quantities were estimated with five tapers and a time-bandwidth product of three using the available toolbox named ‘Chronux’ (http://chronux.org). Time-frequency power spectra (window size of 10 s stepped each 2 s) were calculated for the whole recordings. As the LFPs originating from the same shank were not independent, the median power of these LFPs was calculated to obtain a power estimate for each shank of the polytrode. Since various shanks may record different parts of CA1 with different levels of AAVs infections (Supplementary Fig. 7c), we analysed each shank independently and averaged the data across shank/mouse pairs. The time frequency power spectra were divided into epochs of 30 min. Each one of these epochs was represented by the median over time at each frequency bin.

For the present study, single unit activity (SUA) was also extracted but was simply used as a criterion to select LFPs recorded next to hippocampal neurons. Shortly, unit activity was obtained using the available python-based package named ‘Klusta’ and ‘Klustaviewa’ ^40^. The broadband signal was high-pass filtered at 500Hz and negative peaks were extracted at 2.5 (weak) and 4.0 (strong) standard deviation (STD) thresholds. Spike sorting enabled the isolation of SUAs and only the LFPs recorded at shanks characterized by the presence of SUAs were kept for further analyses.

##### Behavioral testing

Open field tests were performed in healthy adult (3-10 months) animals. Experiments were always done with WT and Lgdel/+ littermates. After surgery was performed, rodents were left to rest for at least three weeks. Mice were habituated to a single experimenter by daily sessions of 5 min handling during the week prior to the behavioural test. Under room lighting conditions (100 lux), locomotor activity was recorded in a black Plexiglas box (43 cm L x 43 cm W x 40 cm H) over which a Camera (Panasonic SD5) was placed. Mice were positioned into the centre of the arena and the distance travelled was measured during 10min using ANY-maze software (Stoelting, UK). To test for the influence of PV neurons on behavioural measures, vehicle (NaCl 0.9%), Neuregulin 1 (NRG1, 50 μg/kg) or clozapine N-oxide (CNO, 2mg/kg) were administered i.p. as previously described ^41,42,43^.

#### Chemical reagents

Carbachol (CCh) was purchased from Sigma-Aldrich. Clozapine-N-oxide (CNO) was purchased from Enzo Life Sciences. A recombinant polypeptide containing the extracellular domain of beta1 Neuregulin 1 (NRG1), able to cross over the brain blood barrier ^44,45^, was purchased from Propsec. Tetrodotoxin (TTX) was purchased from Latoxan. D-2-amino-5-phosphonopentanoic acid (D-APV), 6,7-dinitroquinoxaline-2,3-dione (DNQX), and Gabazine (GBZ) were purchased from Abcam Biochemicals.

#### Statistical analyses

All statistical analyses were done with Matlab, Origin pro 8.0 or the Prism software. We used ANOVA and post-hoc test, Kolmogorov-Smirnov test, Mann-Whitney test and paired Wilcoxon test. All tests were two-sided. For ANOVA, homogeneity of variance was tested using the Mauschly’s sphericity test (subsequent Greenhouse-Geisser or Hunyd-Feldt corrections were applied if sphericity criterion was not met).

## SUPPLEMENTARY FIGURES

**Supplementary Figure 1.**
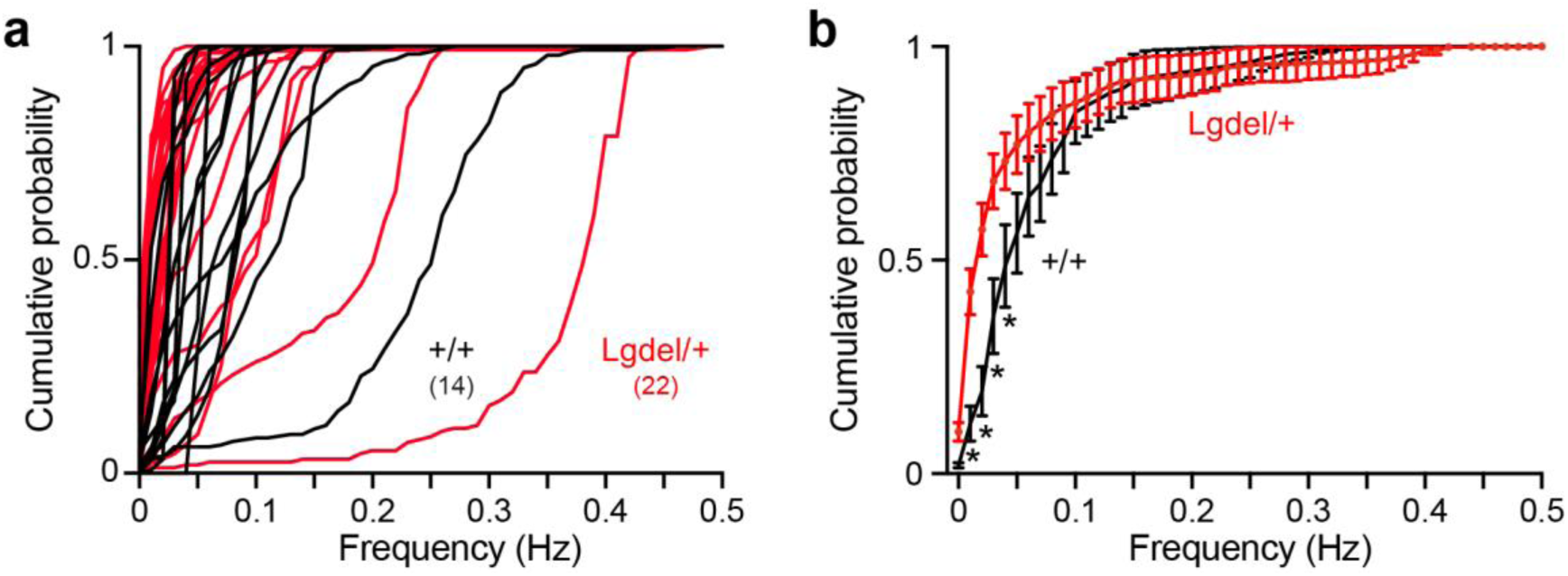
A subpopulation of neurons displays lower calcium transient frequency in Lgdel/+ mice when compared to WT mice. (**a, b**) Cumulative distributions of the frequency of calcium events recorded in Lgdel/+ and WT mice. The distribution is plotted for every slice (**a**) and for the mean of slices from WT or Lgdel/+ animals (**b**). Statistical comparisons were done using 2-Ways repeated measures ANOVA with Bonferroni post-hoc test. \**P* < 0.05. Data are presented as mean ± SEM.

**Supplementary Figure 2.**
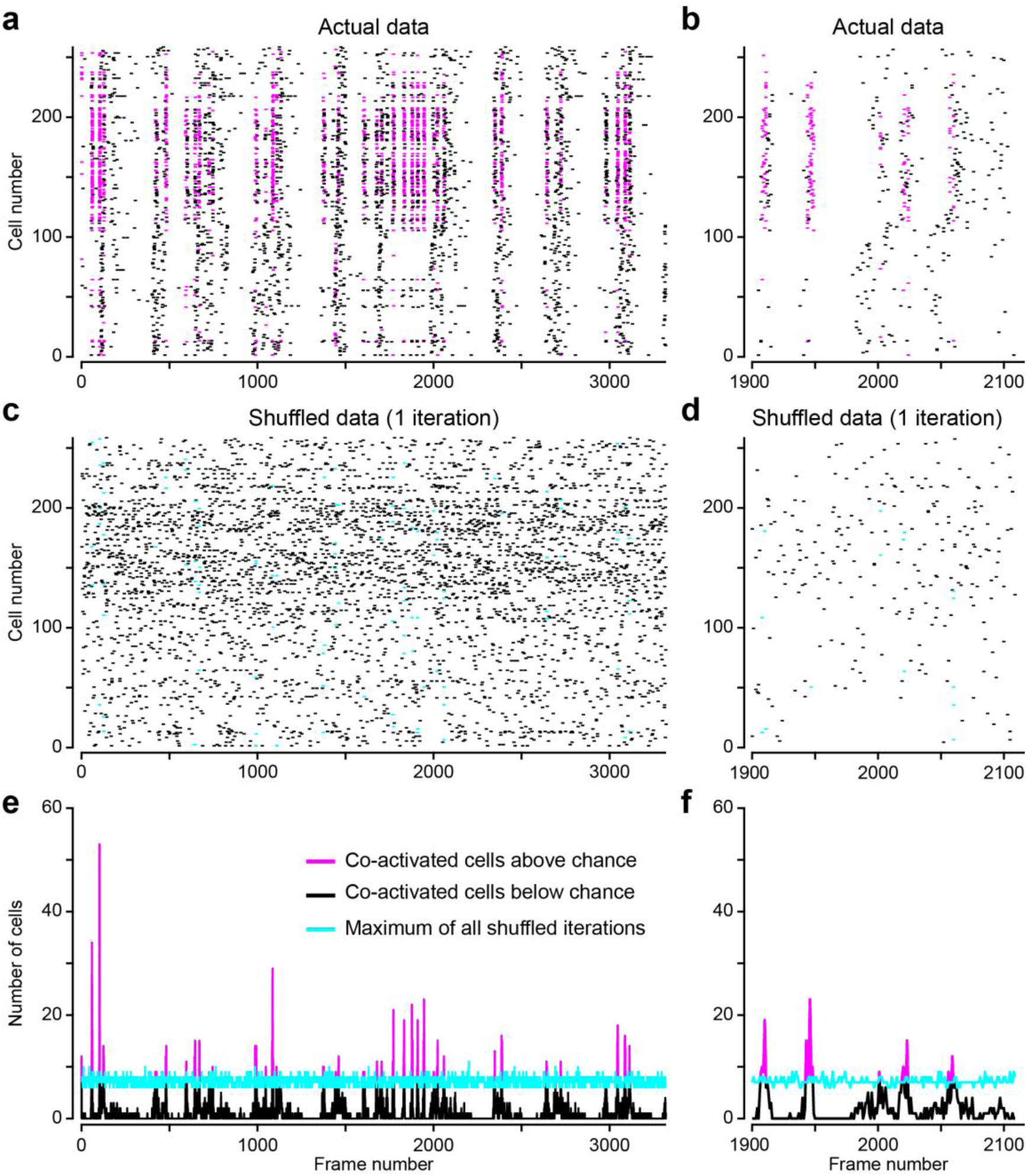
Representative example to illustrate the co-activation analysis of calcium transient onsets. (**a,b**) Raster plots of the calcium transient onsets for the actual data imaged in CA1 neurons of a representative WT slice. (**c,d**) Raster plots of the calcium transient onsets for the same data after one iteration of temporal shuffling (see methods). (**e,f**) A total of 10000 shuffling iterations were made and the iteration with the maximum number of co-activated cells at each frame was chosen as the threshold (magenta). All frames were considered as significant when the number of neurons in the actual data exceeded this threshold and were used to calculate the duration of the co-activations over the whole recordings. The numbers of active cells during these frames were considered as significant co-active neurons (magenta).

**Supplementary Figure 3.**
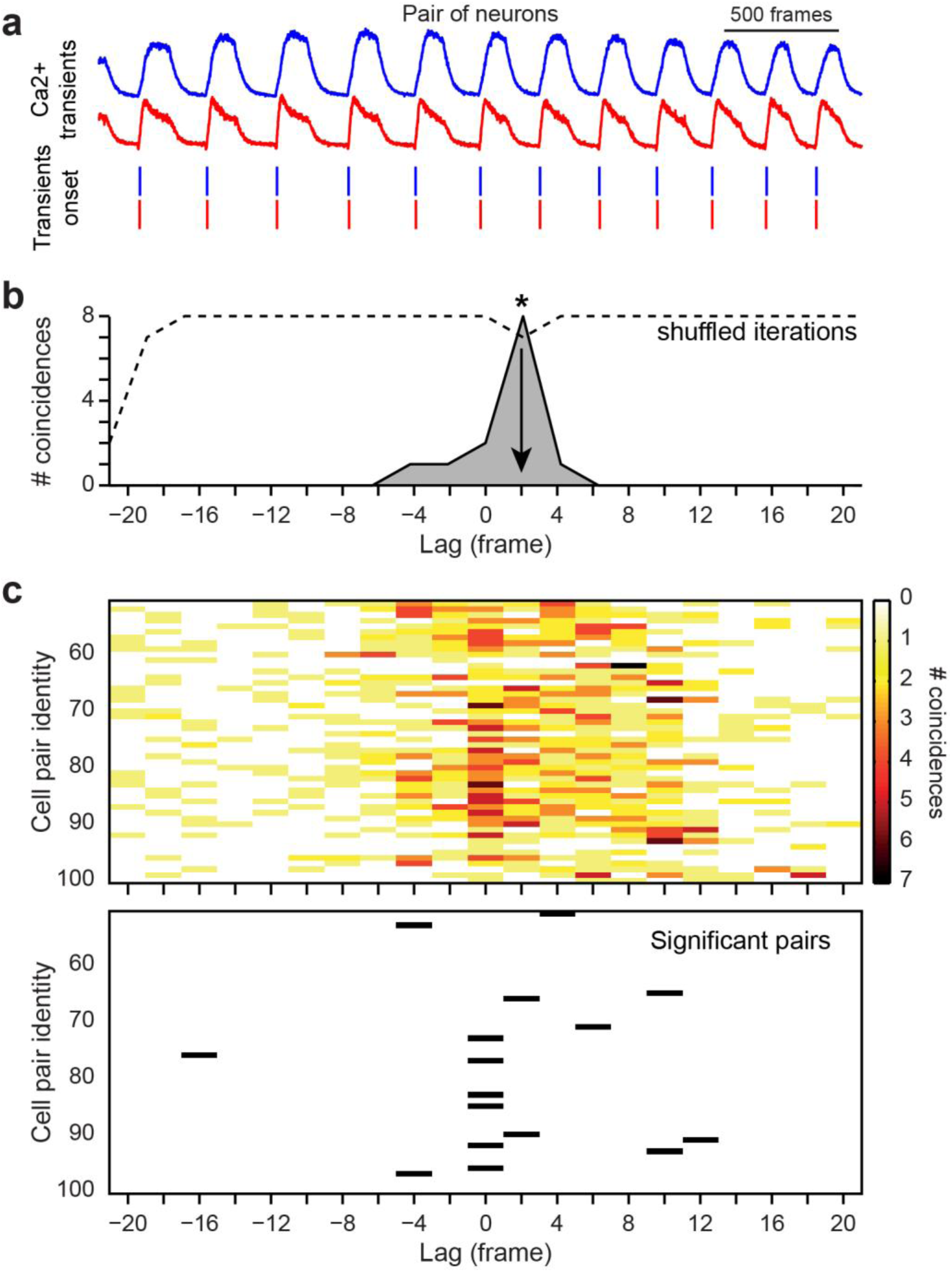
Representative example to illustrate the cross-correlation analysis of paired cells. (**a**) Calcium traces of two neurons (blue and red) and identification of their calcium transient onsets. (**b**) The cross-correlation is presented as the number of coincidences (shaded area) as a function of frame lags. A temporal shuffling procedure was used to estimate the significance threshold (dashed line). (**c**) Each time the number of coincidences exceeded the threshold, the corresponding frame lag (lower panel) was considered as significant. Data is shown for few pairs of neurons.

**Supplementary Figure 4.**
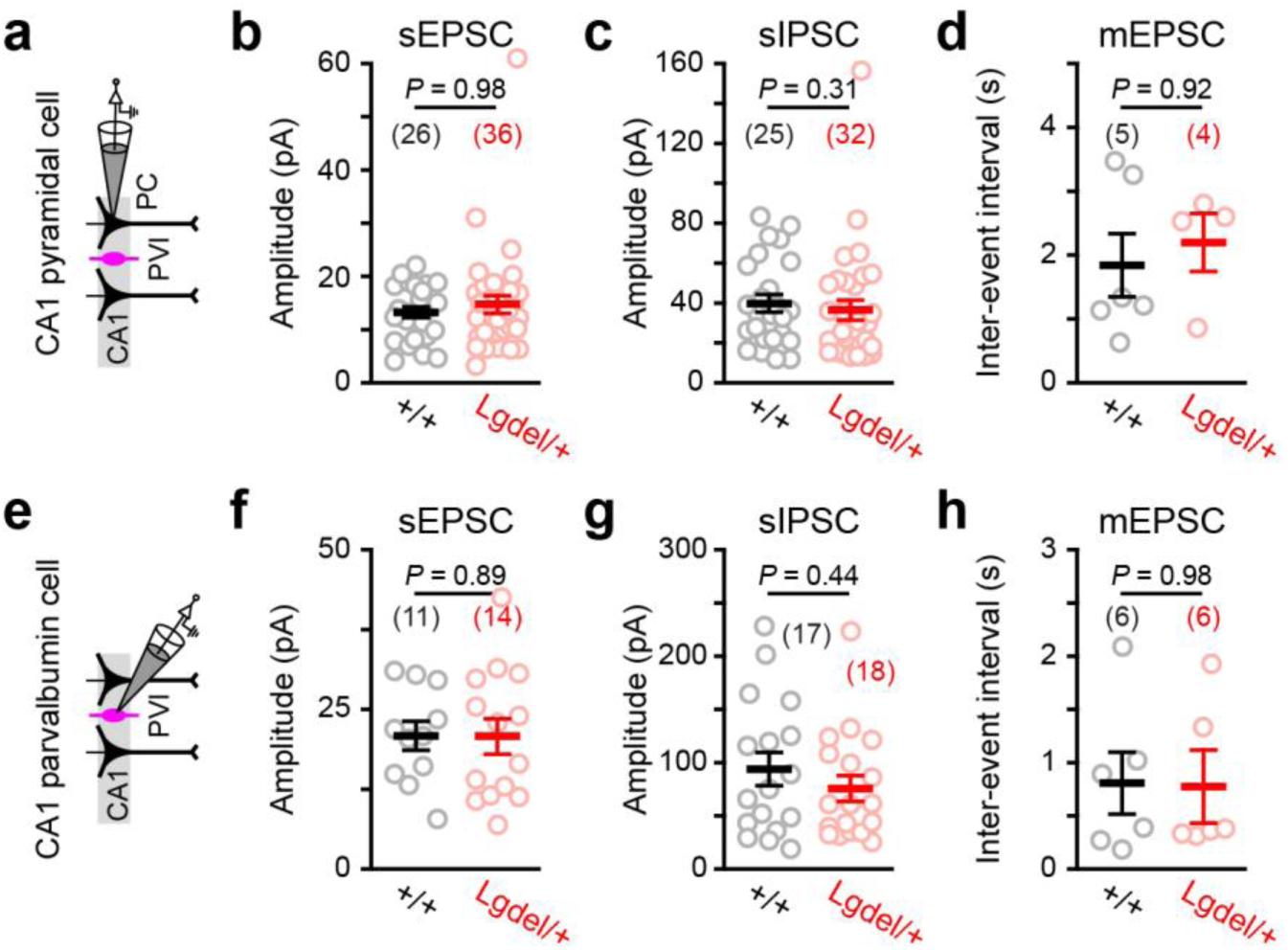
Synaptic parameters recorded in PCs and PVIs from WT and Lgdel/+ mice. (**a-d**) Patch-clamp recordings of CA1 PCs from WT and Lgdel/+ mice. Summary graphs of the mean sEPSC (**b**) and mean sIPSC (**c**) amplitude. (**d**) Summary graph of the mean mIPSC IEI. (**e-h**) Patch-clamp recordings of CA1 PVIs from WT and Lgdel/+ mice. Summary graphs of the mean sEPSC (**F**) and mean sIPSC (**g**) amplitude. (**h**) Summary graph of the mean mIPSC IEI. Data are presented as mean ± SEM (each circle represents a cell, number of cells indicated in parenthesis; Mann-Whitney test).

**Supplementary Figure 5.**
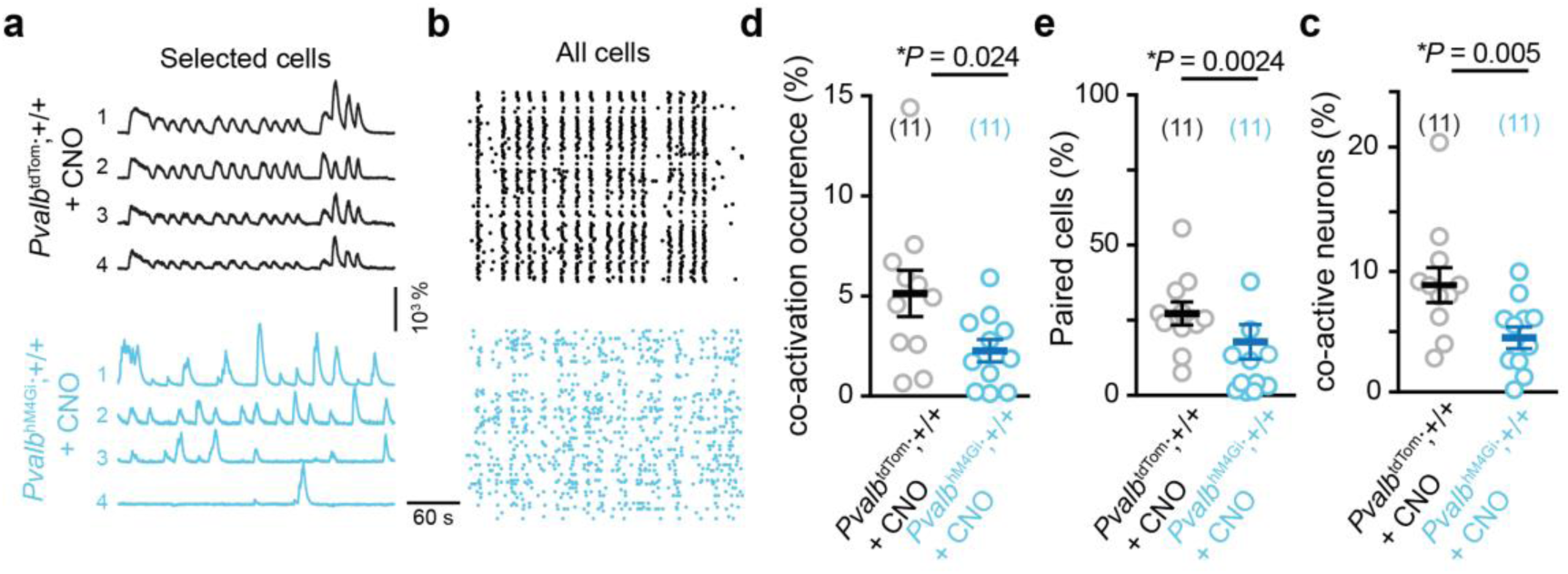
Chemogenetic inhibition of PVI desynchronizes CA1 assemblies in WT mice. (**a**) Examples of Ca^2+^ sweeps recorded from four selected neurons in slices of *Pvalb*^cre/+^;+/+ mice infected with AAVs enabling the conditional expression of either tdTomato (*Pvalb*^tdTom^;+/+) or the inhibitory DREADD hM4Gi (*Pvalb*^hM4Gi^;+/+) after clozapine-N-oxide (CNO, 1 μM) bath application. (**b**) Raster plots of Ca^2+^ transient onsets extracted from all neurons of a recorded field. (**c-d**) Percentage of co-active neurons above random co-activation (**c**) and occurrence of these co-activations (**d**). (**e**) Percentage of correlated pairs of neurons above chance. Statistical comparisons were done with Mann-Whitney test; \**P* < 0.05. Data are presented as mean ± SEM.

**Supplementary Figure 6.**
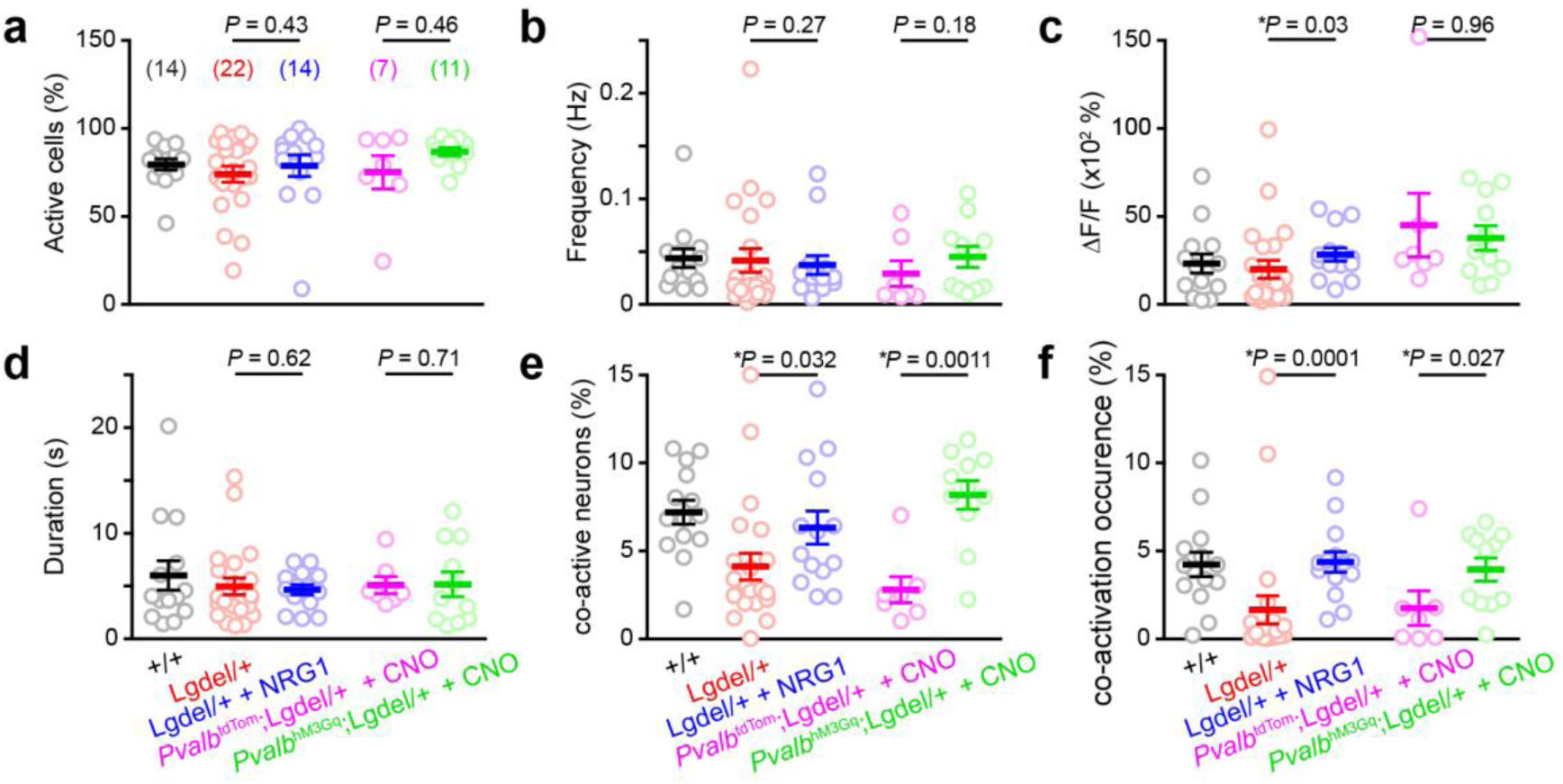
Effect of PVI activation on various network dynamics parameters in Lgdel/+ mice. (**a**) Proportion of neurons displaying Ca^2+^ transients. (**b-d**) Frequency (**b**), amplitude (**c**) and duration (**d**) of Ca^2+^ transients recorded in CA1 neurons. (**e**) Percentage of co-active neurons above random co-activation. (**f**) Occurrence of the co-activations. Statistical comparisons were done with Mann-Whitney test; \**P* < 0.05. Data are presented as mean ± SEM.

**Supplementary Figure 7.**
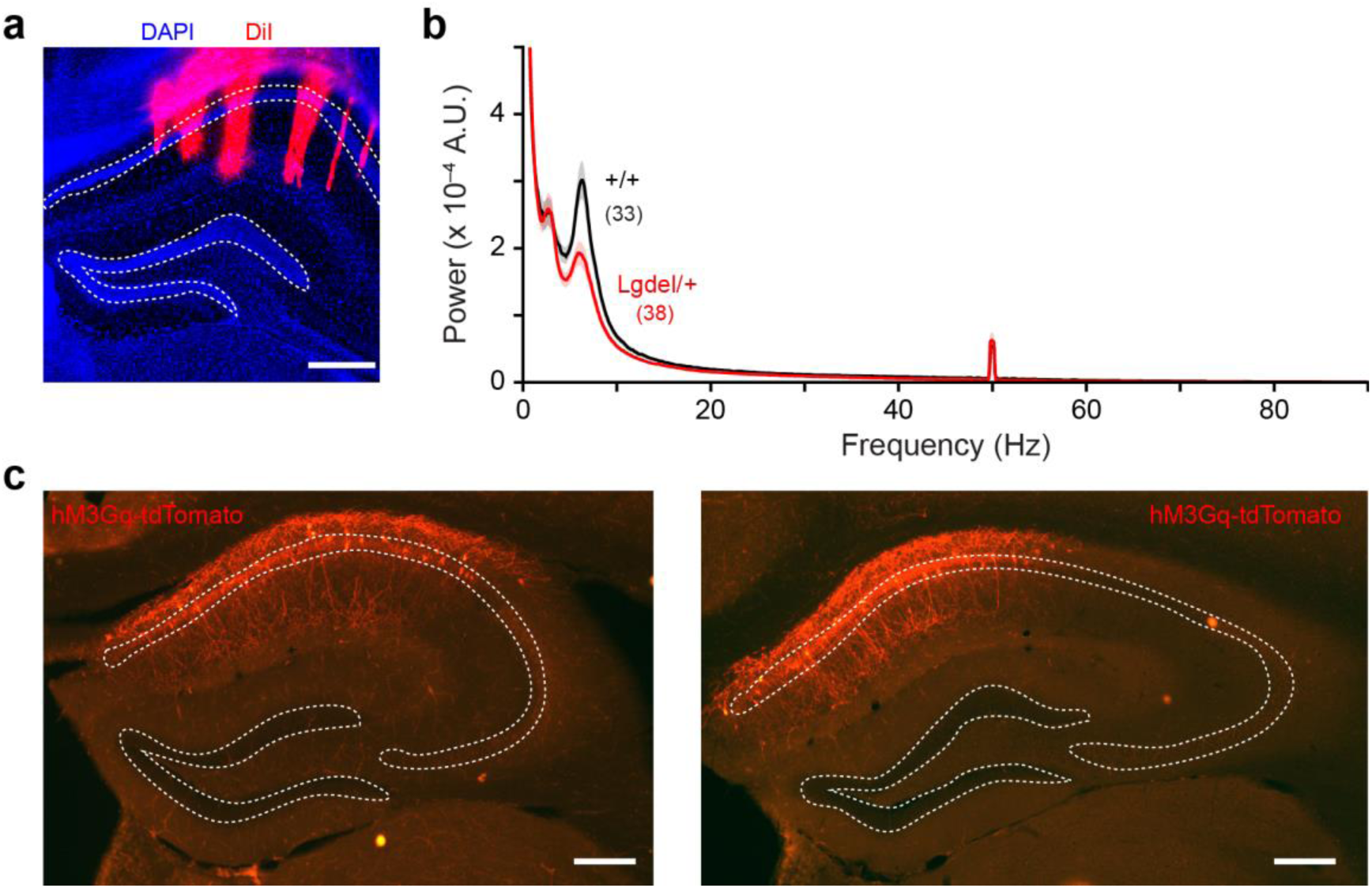
*In vivo* LFP recordings using polytrodes and DREADD infection in the hippocampus. (**a**) Photograph of the track resulting from the implantation in dCA1 of the silicon polytrodes coated with DiI. For each shank, the recording sites close the CA1 layer were used for LFP spectral analysis (see methods). (**b**) Distributions of LFP power spectra for WT and Lgdel/+ (mean ± SEM; number in parentheses indicate the numbers of mouse/shank pairs). (**C**) Photographs showing the AAV-mediated expression of hM3Gq-tdTomato in dCA1 PVI from two different mice.

**Supplementary Figure 8.**
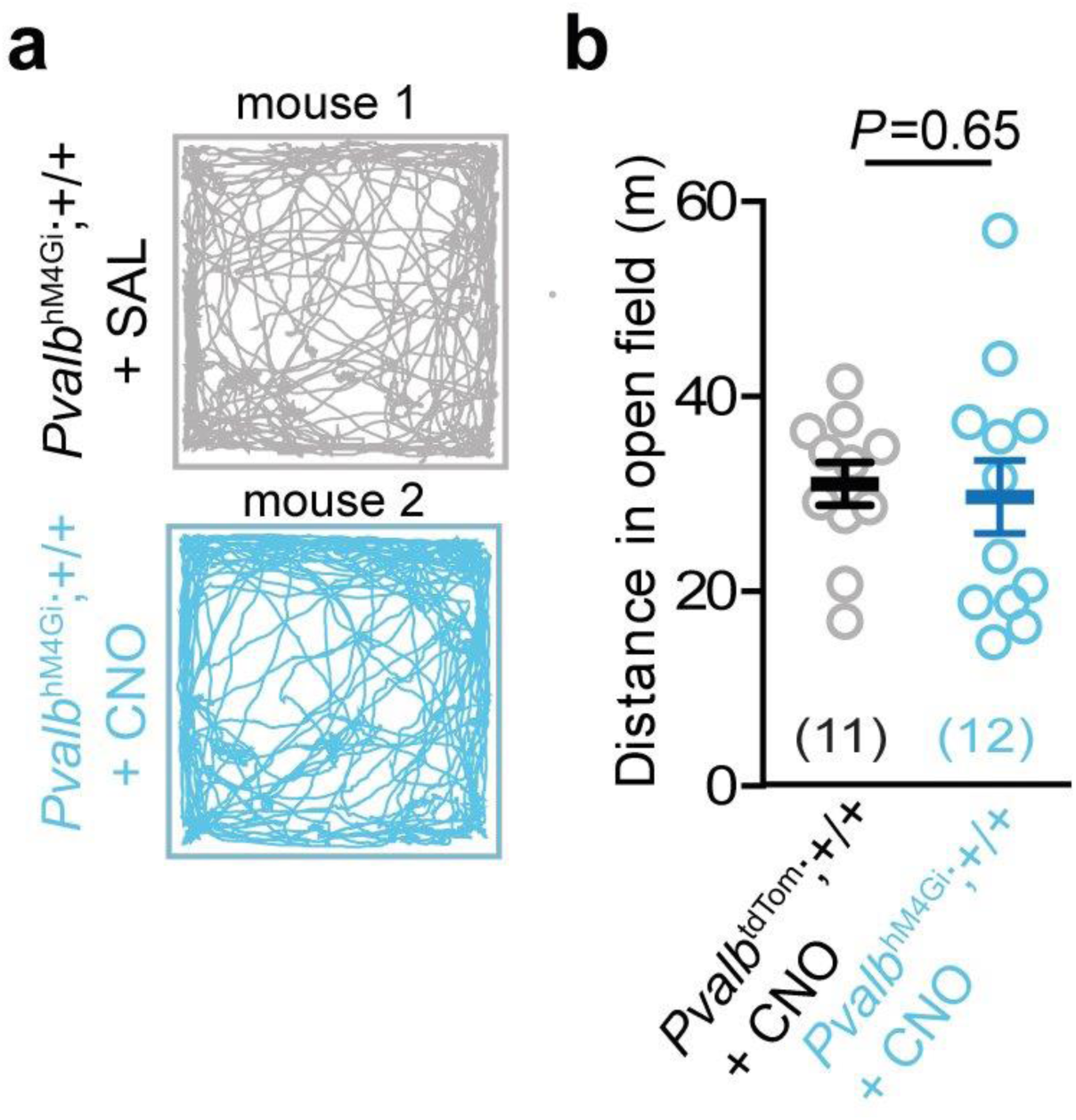
Effect of the chemogenetic inhibition of PVI on behaviour in the Open Field Test. (**a**) Locomotor tracks of different transgenic mice and treatments during 10 min in an open field. (**b**) Summary graph of the distance travelled in the open field. Statistical comparisons were done with Mann-Whitney test; \**P* < 0.05. Data are presented as mean ± SEM.

